# NetQuilt: Deep Multispecies Network-based Protein Function Prediction using Homology-informed Network Similarity

**DOI:** 10.1101/2020.07.30.227611

**Authors:** Meet Barot, Vladimir Gligorijević, Kyunghyun Cho, Richard Bonneau

## Abstract

Transferring knowledge between species is challenging: different species contain distinct proteomes and cellular architectures, which cause their proteins to carry out different functions via different interaction networks. Many approaches to proteome and biological network functional annotation use sequence similarity to transfer knowledge between species. These similarity-based approaches cannot produce accurate predictions for proteins without homologues of known function, as many functions require cellular or organismal context for meaningful function prediction. In order to supply this context, network-based methods use protein-protein interaction (PPI) networks as a source of information for inferring protein function and have demonstrated promising results in function prediction. However, the majority of these methods are tied to a network for a single species, and many species lack biological networks. In this work, we integrate sequence and network information across multiple species by applying an IsoRank-derived network alignment algorithm to create a meta-network profile of the proteins of multiple species. We then use this integrated multispecies meta-network as input features to train a maxout neural network with Gene Ontology terms as target labels. Our multispecies approach takes advantage of more training examples, and more diverse examples from multiple organisms, and consequently leads to significant improvements in function prediction performance. Further, we evaluate our approach in a setting in which an organism’s PPI network is left out, using other organisms’ network information and sequence homology in order to make predictions for the left-out organism, to simulate cases in which a newly sequenced species has no network information available.

## 1. Introduction

Sequences have been the primary source of information protein function prediction, mainly because of their abundance and the ease with which many models can incorporate large amounts of sequence data. However, for function prediction, sequence information fails to give the context of a protein in an organism; this context can be highly relevant in determining the protein’s function. Protein interaction networks, on the other hand, offer a way to understand how proteins function in cellular pathways, and thus have been a powerful source of information for inferring the functions of unannotated proteins [1, 2, 3, 4, 5].

In community benchmarks, such as the Critical Assessment of Functional Annotation (CAFA), the best-performing methods rely on multiple complementary data sources — protein sequence, structure, and network information — in order to make more accurate predictions [6, 7, 8]. There are many reviews of protein function prediction methods in general [9, 10, 8, 11]. Most previous network-based approaches integrate different types of networks containing complementary information to achieve state-of-the-art performance[12, 4, 3], but are limited to training on and making predictions for a single organism’s proteins. Methods for sequence and structure-based function prediction are numerous [13, 14, 15]; these methods are inherently able to predict functions for proteins of multiple organisms, and can have certain other advantages such as region specificity for predictions [15, 16]. A remaining challenge is using the vast amounts of network information from multiple species in a single model.

Our method, NetQuilt, accomplishes several important goals in function prediction. First, NetQuilt allows for the integration of sequences and networks, which allows the limited knowledge of the homology between proteins to be supplemented by knowledge of the network topology, and vice versa – incomplete protein-protein interaction networks are supplemented by homology. NetQuilt also creates protein features that are not tied to single species and that include evolutionary and functional information. As a result of the increased training examples in the multispecies setting compared to methods considering only single species, rarer Gene Ontology (GO)[17] terms are able to be trained on. The much larger set of training examples also serves to improve prediction on more abundant terms. Most importantly, our method enables network-based function prediction even for species for which knowledge of their protein interaction networks is limited. We demonstrate the achievement of these goals in several settings. We compare the quality of protein features of a single organism in a single-species versus a multispecies setting. We show that multispecies features are more indicative of a protein’s function than single-species features. We also test the model’s ability to predict functions of a species whose entire PPI network is missing, with the model trained on all other species in the set being considered, in an approach termed “leave one species out” (LOSO). We demonstrate that our model is capable of using information from other species to correctly infer functions of the missing species.

## 2. Related work

### 2.1. Graph node classification methods

Protein function prediction using PPI networks is a node classification problem, the methods for which can be categorized into two groups: label-propagation methods, and classifiers trained on graph features. Label propagation methods propagate labels from labeled nodes to unlabeled nodes via random walks; this strategy is used to predict protein function in a method called GeneMANIA [3]. The category of classifiers trained on graph features can be split further into two categories: those that manually engineer features from the network data, or those methods that learn network embeddings of nodes in order to be used in a classifier. The manually engineered graph features can be based on graph measures such as node degree, neighborhood size within some number of steps, number of shortest paths, etc. Other features that can be constructed over nodes include graphlets [18], and random walk profiles of nodes within their graph, which have been extended and applied to heterogeneous and multiplex biological networks [19, 20]. Network embedding has been extensively used in protein functional analysis and includes methods based on matrix factorization [4], graph kernels [21] and deep learning [12, 22, 23]. A comprehensive review of network embedding in computational biology compared to other types of network-based algorithms for several applications can be found in [24], and reviews of network representation learning methods in general can be found in [25] and [26].

### 2.2. Single-species network-based methods

Our previous study [12] introduced a method called deepNF (deep Network Fusion), which involves using a multimodal autoencoder to create embeddings of nodes from different types of protein-protein interaction networks of an organism. These embeddings are then used to train support vector machines (SVM) to predict GO terms. This method outperformed other methods using different types of interaction networks to predict function, including Mashup[4] and GeneMANIA[3], all of which had access to six STRING network types {‘experimental’, ‘coexpression’, ‘coocurrence’, ‘neighborhood’, ‘fusion’, and ‘database’}. This work demonstrated that multimodal autoencoder neural networks could effectively extract functionally informative features from graphs with multiple edge types. Another method, STRING2GO, uses maxout neural networks in order to create functional representations of proteins from protein interaction networks of a single species [22]. The max-out network is trained to predict GO terms from Mashup or Node2Vec [27] node embeddings, and the representations of each protein is taken from the layer before the output predictions. These representations are then used to train SVMs to predict GO terms. The authors show that these representations are able to outperform the original Mashup and Node2Vec embeddings of PPI networks when used to train SVMs for the function prediction task. In [23], an unsupervised neural network is used to learn embeddings from a tissue-specific multi-layer PPI graph. These task-independent embeddings are then used to predict multi-cellular function.

However, these methods are limited to using information from single organisms for prediction, because they operate on a feature space common only to proteins of that organism. A better approach would be to take into account information from proteins of many different organisms at once in order to take advantage of large-scale training sets.

### 2.3. Multispecies methods

A few methods make use of information from protein interaction networks of multiple species. One such method is NetGO, an ensemble learning-to-rank method that combines six component methods, one of which is a k-nearest-neighbors method that uses PPI networks of multiple species [28]. One drawback to this method is that it is unable to use the homology information in any way beyond direct transfer of annotation between homologues. Ideally, a protein function prediction method should be able to use homology information to supplement network information even on proteins whose sequences are not similar to the training set protein sequences. Another method, MUNK, is a kernel-based method that produces functional embeddings used for predicting synthetic lethality for pairs of proteins of multiple species [21]; they additionally demonstrate that proteins close in this embedding space are similar in function. The key idea of their approach is that proteins from different species are embedded in the same vector space using graph kernels with landmark proteins in the networks of the two species that perform the same functions.

### 2.4. Global network alignment and IsoRank

The problem of network alignment is to find topological and functional similarities between nodes of different networks. Local network alignment algorithms aim to find subgraphs which are conserved between input networks, while the goal of global network alignment algorithms is to find mappings of all nodes between the input networks. Most network alignment methods focus on this latter goal [29, 30, 31, 32, 33, 34, 35]. IsoRank[29] is a global network alignment algorithm used to align multiple PPI networks. This is done in two stages: first by solving an eigenvalue problem across all pairs of input networks to obtain protein similarity scores, and then by using k-partite matching to obtain the final alignment of all organisms, giving sets of functional orthologs across species. IsoRankN[30] was developed as an improvement to the alignment extraction portion of IsoRank in which instead of k-partite matching, spectral clustering was applied to the metagraph of all organisms’ proteins induced by the similarity scores given by the eigenvalue problem. More recent global network alignment algorithms include L-GRAAL [31], which uses a graphlet similarity-scoring function used with a search heuristic based on Lagrangian relaxation, and GHOST, whose key step uses a signature of nodes based on the spectrum of the normalized Laplacian of local subgraphs; this signature is then used to measure topological similarity of networks [34]. Fuse [35] is another network alignment method consisting of two steps. The first step calculates functional similarity between proteins using a weighted sum of scores from a nonnegative matrix tri-factorization of all considered PPI networks and sequence similarity. The second step constructs an edge-weighted k-partite graph (where k is the number of PPI networks) from these similarities and then obtains the one-to-one network alignment using an approximate maximum weight k-partite matching solver. A comprehensive review of biological network alignment can be found in [36]. Other algorithms for network alignment include those that focus on finding small network region similarities conserved among networks, unconstrained by the assumption of one-to-one mapping of nodes. These algorithms fall into the local network alignment category. A comparison study of local and global network alignment methods can be found in [37], where it was found that network topology has additional biological knowledge compared to sequence data; additionally, global and local network alignment methods may give complementary information for protein function prediction.

In this study, we use the first step of IsoRank to integrate sequence homology information with PPI network information to generate functionally-informative similarity scores between species as well as within species themselves. We use these similarity scores for every protein as its feature representation to enable the training of a neural network with proteins coming from many different organisms’ PPI networks in the same input space.

## 3. Methods

In this section, we outline the components of our method, NetQuilt. These components are the global network alignment algorithm for creating both intranetwork (within-species) and internetwork (between proteins in different species) node-similarity profiles, and the maxout neural network, which uses the concatenated aligned-network vectors to predict Gene Ontology (GO) terms. See Fig. 1 for an overview of the procedure.

**Figure 1:**
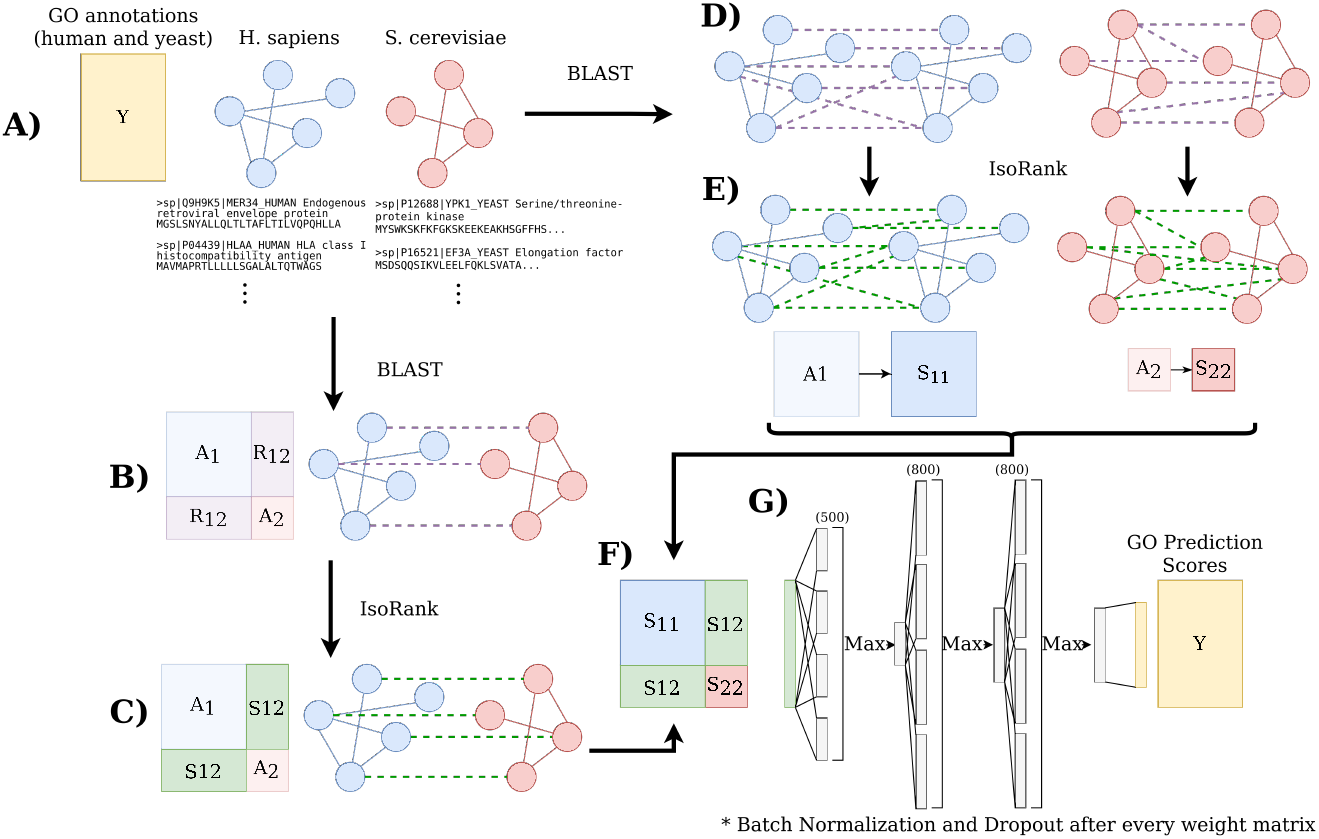
Overview of our method for running on two organisms (human and yeast). **A)** For each taxonomy ID, download network, annotation and sequence files from the STRING-db static website (version 11). **B)** Use BLAST to create sequence identity links between proteins of pairs of different species. **C)** Compute IsoRank scores between proteins of different species, using BLAST sequence identity values and the organisms’ networks to create a combination of network and homology information. **D)** Use BLAST to create sequence identity links among proteins of each individual species. **E)** Compute IsoRank alignment scores between proteins of the same species, creating denser matrices *S*_11_ and *S*_22_ from weighted adjacency matrices *A*_11_ and *A*_22_ and sequence identity matrices *R*_11_ and *R*_22_. **F)** Concatenate all IsoRank matrices between all species to make the full S matrix. **G)** Train maxout neural network with the S matrix as features and the annotation matrix as labels.

### 3.1. Creating multispecies similarity profiles with IsoRank

Consider a set of *N_org_* undirected graphs, where each graph is a proteinprotein interaction network of a different organism. The graphs each have a set of nodes representing proteins for each organism, and a set of edges representing the interactions between these proteins. The graphs are represented by adjacency matrices {**A**^(1)^, **A**^(2)^,…, **A**^(Norg)^}. See Figures 3, 2, and Supplemental Tables 1 and 2 for statistics on the networks used in this study. Consider further that we have a set {**R**_1;1_, **R**_1,2_, **R**_1,3_,…, **R**_1,*Norg*_, **R**_2,2_, **R**_2,3_, **R**_2,4_ …, **R***_Norg,Norg_*} of edges of another type, between all proteins of all species. Our method computes profiles of the nodes in all species’ networks, creating a shared feature space for all proteins, which we then use to train a maxout neural network to predict protein function. We first compute similarity scores between proteins of different species in a way derived from the IsoRank method of multispecies network alignment [29]. The scores are given by the following recurrence equation:

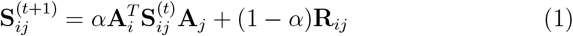

where:

- 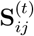 is the similarity matrix between networks (species) *i* and *j* after *t* steps of diffusion;
- **R**_*ij*_[*k,l*] = − log(*eval*[*k,l*]) is the blast e-value similarity between protein *k* in network (species) *i* and protein *l* in network (species) *j*; and
- **A**_*i*_, **A**_*j*_ are the row-normalized adjacency matrices of networks (species) *i* and *j*.

**Figure 2:**
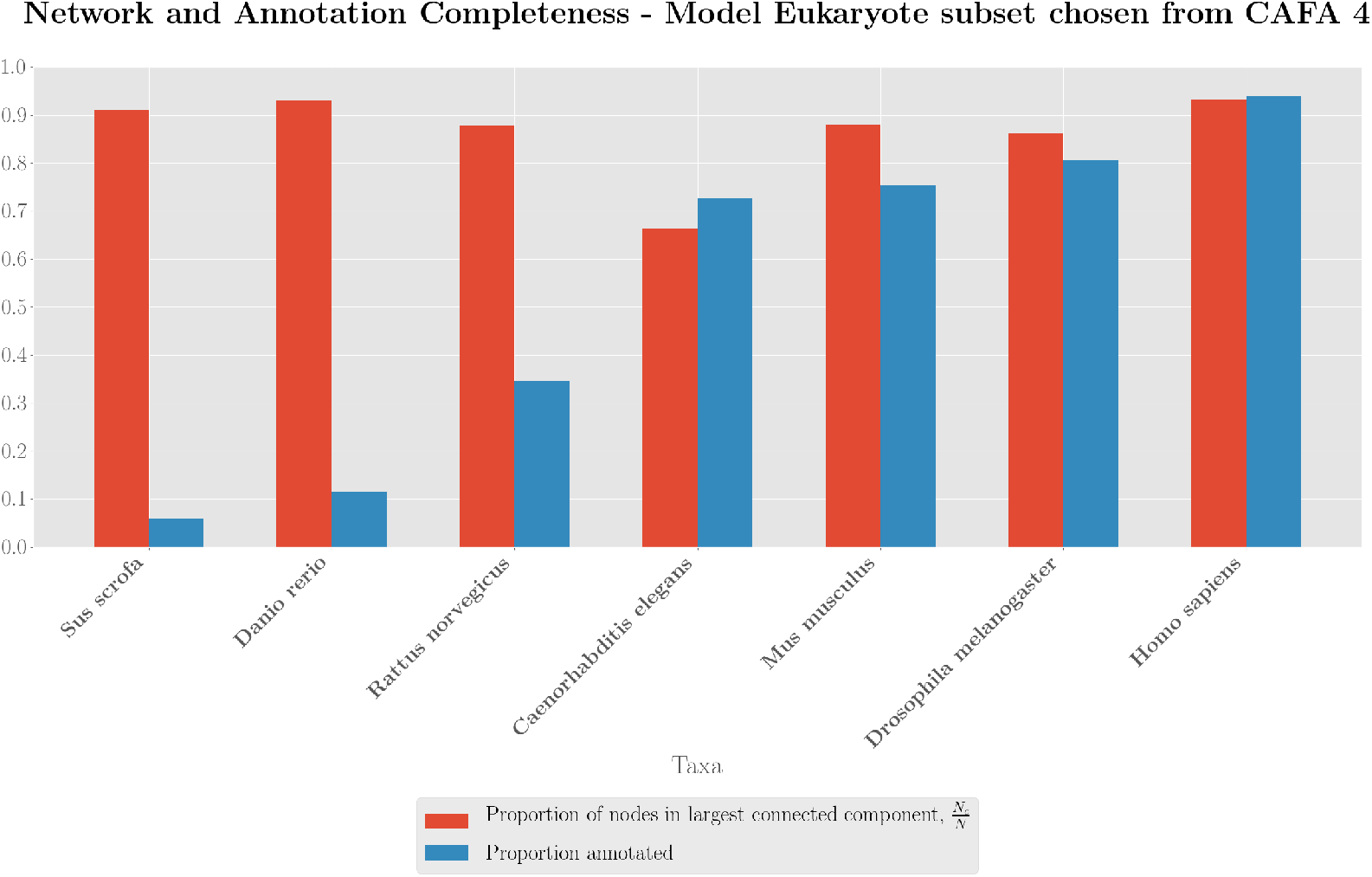
Proportions of STRING proteins of a given Eukaryote taxonomy ID that are annotated in at least one branch of the Gene Ontology with any evidence code and proportions that are in the STRING experimental network’s largest connected component, sorted by the sum of the two proportions of each species.

**Figure 3:**
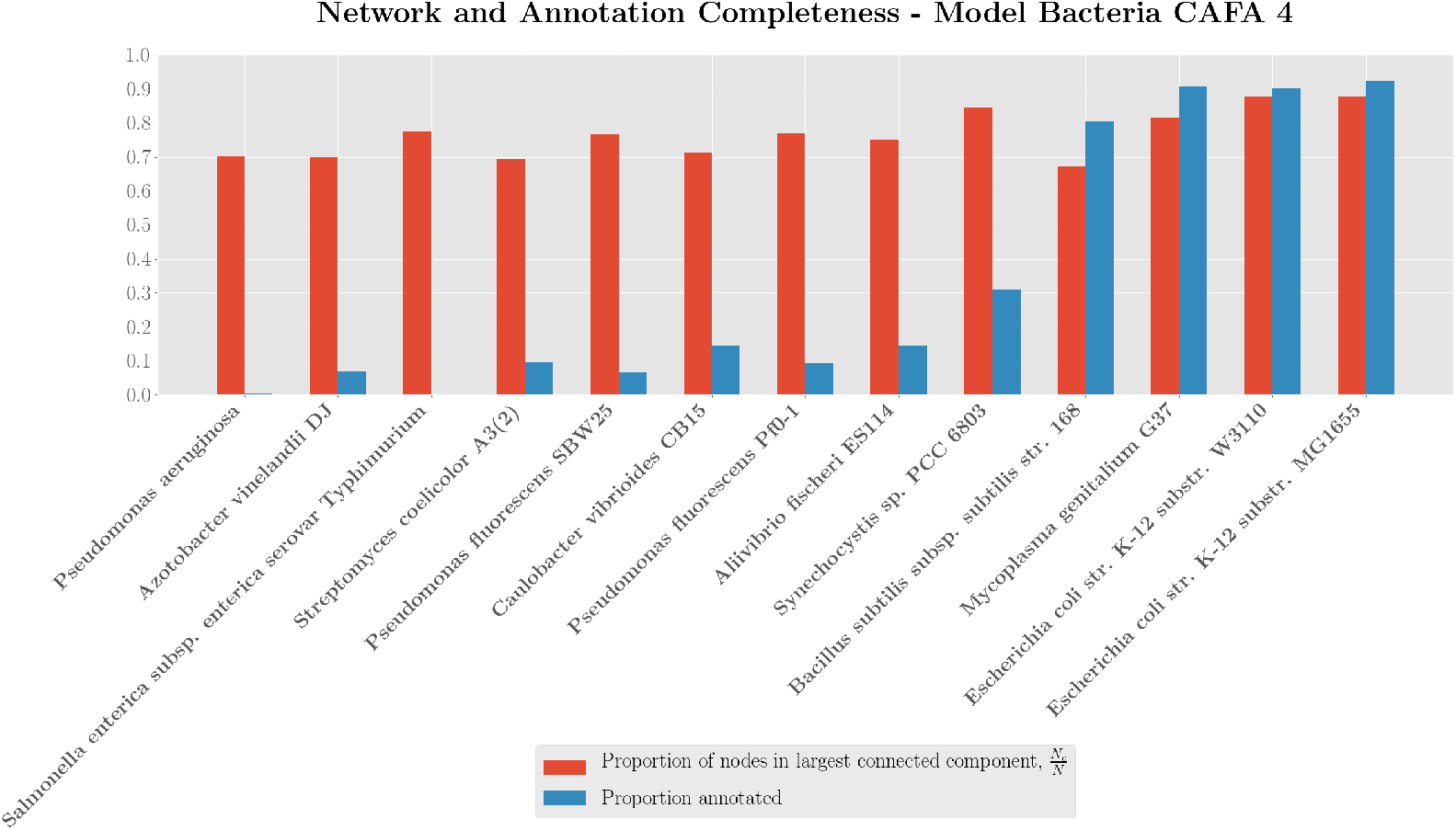
Proportions of STRING proteins of a given Bacteria taxonomy ID that are annotated in at least one branch of the Gene Ontology with any evidence code and proportions that are in the STRING experimental network’s largest connected component, sorted by the sum of the two proportions of each species.

Starting with **S**^(0)^ = **I**^*n*_*i*_×*n*_*j*_^, we iterate this calculation (Equation 1) until convergence with respect to the norm of the difference between the previous matrix 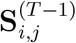 and the current matrix 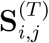. We then calculate IsoRank similarity scores between proteins *within* each species. This computes ‘‘alignment” scores between a network and itself, integrating (normalizing) sequence homology scores computed using BLAST and protein-protein interactions.

We can now construct a large symmetric matrix **S** in which the IsoRank similarity matrices of all species with themselves are placed along the diagonal, resulting in a block-diagonal matrix. Next, each interspecies protein similarity matrix **S_ij_** is placed on the off-diagonal, comprising the submatrix with row indices of the proteins of species *i* and column indices of the proteins of species j. Refer to steps B, C, D, E & F in Fig. 1 for a visual description of this matrix construction. **S** now contains the information from all the individual protein interaction networks as well as the links between them, integrated with sequence-similarity information. We finally use this matrix as input to a maxout neural network, with each row of the matrix **S** being used as a single training sample.

### 3.2. Using maxout neural networks to predict protein function from aligned meta-network features

Maxout neural networks, introduced in [38], are neural networks whose layers have the maxout activation function. The maxout activation of a layer is the element-wise maximum of a set of affine transformations to the input of that layer. More explicitly, a maxout layer’s ith output value 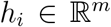 given an input 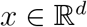 is defined as:

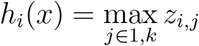

where *z_ij_* = *x^T^W_:i,j_* + *b_i;j_* is the ith element of the jth affine transformation of the input vector with learned parameters 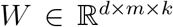 and 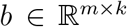. Maxout activation functions are able to approximate arbitrary convex functions, and therefore enable the neural network to learn not only relationships between hidden units but also the activation functions themselves. This provides additional flexibility, which enables the neural network to learn features that are more specifically tailored to a prediction task.

The architectures for our models are listed in Table 1 (see also part G in Fig. 1). To avoid overfitting, we use early stopping with the criterion of improving AUPR calculated over a validation set consisting of 20% of the training data, with patience 30 (i.e., if the AUPR score does not improve in 30 consecutive epochs, the training is stopped). The architectures were chosen using cross-validation performance on datasets for eukaryotes and bacteria using the previous version of STRING (v10.5) [39] for annotations and network information. The hyperparameter search started with an architecture based on [22], with three rounds of random search, trying 1% of possible models each round. Empirically, maxout neural networks performed better than neural networks with sigmoid or ReLU activation functions for this task. Other benefits of maxout neural networks include fast gradient computations relative to other activation functions, e.g. sigmoid, and fewer choices of hyperparameters, since the activation function is learned. The search space was restricted after each round by removing possible values of hyperparameters that were correlated with lower performance. The models were implemented using Keras [40]; the code is freely available at https://github.com/nowittynamesleft/NetQuilt.

**Table 1:** Model architectures for Eukaryote and Bacteria datasets (see Section 4.1 for a description of these datasets). “Maxout units” refers to the number of separate weight matrices for a given layer; the element-wise max is computed over the product of the weight matrices with the outputs of the previous layer.

| **Hyperparameters** | **Bacteria** | **Eukaryotes** |
| --- | --- | --- |
| Hidden Layer Dimensions | [500, 800, 800] | [500, 800, 800] |
| Maxout Units | 3 | 4 |
| Dropout | 0.2 | 0.2 |
| Batch Normalization[41] | True | True |
| Learning Rate | 0.01 | 0.01 |
| Batch Size | 16 | 32 |
| Max Number of Epochs | 100 | 300 |
| Optimizer | AdaGrad[42] | AdaGrad |

## 4. Experiments

### 4.1. Datasets

We conduct our analyses on both a collection of eukaryote networks and a separate collection of bacteria networks. Each dataset consists of STRING PPI networks, of which we use only the “experimental” category for our method, and Gene Ontology annotations of each organism retrieved from STRING version 11 [39]. The statistics on the organisms we include in our study are given in Figures 2 and 3, which show the networks’ largest connected component ratios and the annotation percentages of proteins present in STRING. The numbers of nodes and edges, for bacteria and eukaryotes, are shown in Supplemental Tables 1 and 2 respectively. In order to select the value of the *α* parameter for our experiments for each set, we tested several values in a single-species cross-validation setting (see Supplemental Figures 3, 4, and 5 for the results of the search). The chosen organisms come from the set of organisms that were evaluated in CAFA 4. For the bacteria, all of the organisms from CAFA 4 were used in our pipeline; for the eukaryotes, we selected a subset to conserve memory when training our models (C. elegans, D. melanogaster, D. rerio, H. sapiens, S. scrofa, M. musculus, and R. norvegicus). We use GO terms that cover between 0.5% - 5% of the species’ proteins in its PPI network, and remove proteins without annotations of these GO terms from training and evaluation sets.

**Figure 4:**
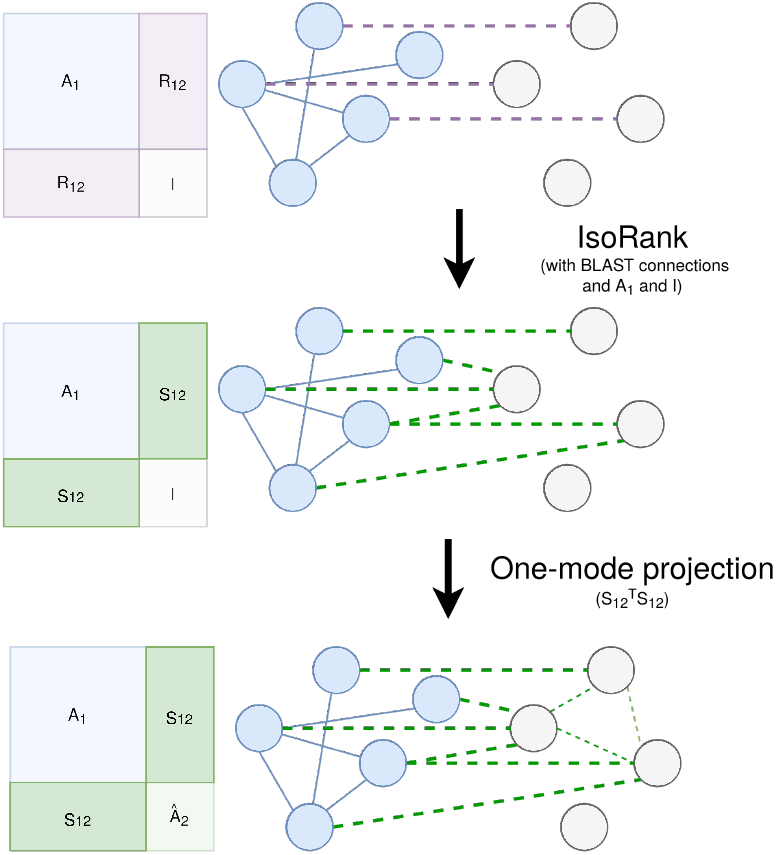
Procedure for predicting a network to be used in the leave-one-species-out validation setting, where we assume no knowledge of the PPI network for one organism. First, BLAST connections (represented as purple dashed lines) between the proteins of the known network and the left-out network are created. IsoRank is then run for the interspecies matrix, using the known network *A*_1_ and the left-out network given by the identity matrix I, giving the IsoRank connections *S*_12_ depicted by the large green dashed lines. We finally obtain a predicted network by taking the one-mode projection of the IsoRank connections: 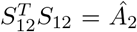. In the case of multiple known organisms, we simply take the average of all organisms’ one-mode projections with the left-out organism.

**Figure 5:**
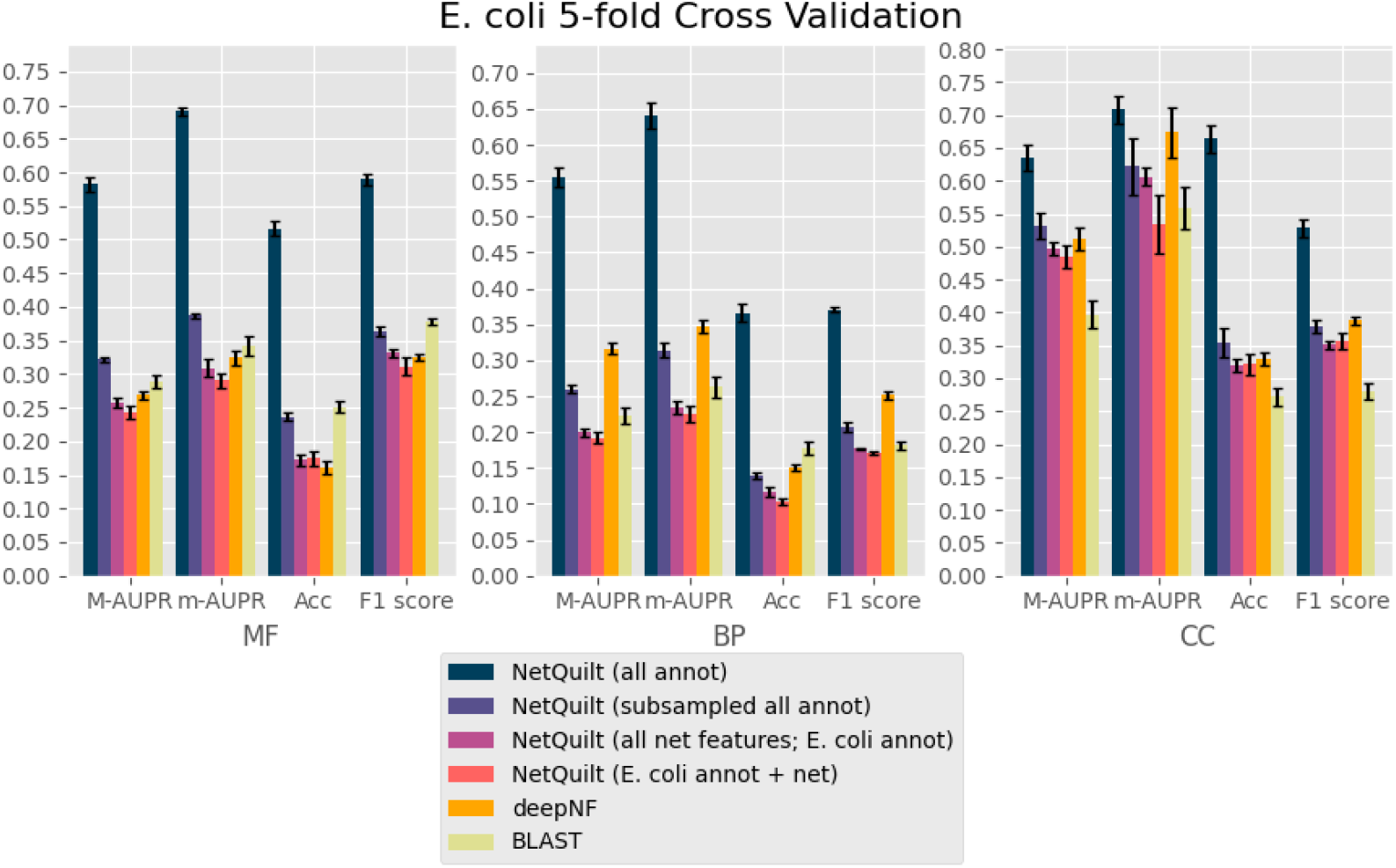
Performance comparison of NetQuilt method with baselines. Methods shown: NetQuilt trained on model bacteria annotations; NetQuilt trained on subsampled model bacteria annotations; NetQuilt trained only on E. coli str. K-12 substr. MG1655 examples; single-species NetQuilt (taking only the E. coli IsoRank matrix and annotations); deepNF (single-species, but integrating 6 STRING network types); and CAFA BLAST annotation transfer method using all selected bacteria annotations.

### 4.2. Cross-validation

In our first set of evaluations, in order to compare with single-species methods, we perform cross-validation on a single test species at a time. The performance is averaged over 5 repetitions with 20% of data used as the test set. We train our models on GO term annotations of any evidence code[17], but evaluate our predictions with annotations of the evidence codes EXP, IDA, IPI, IMP, IGI, IEP, TAS and IC, as previously used in CAFA papers [6]. Since, realistically, our method has access to more training examples than the single-species methods, we include three benchmark versions of our method:

1. NetQuilt trained on a subsampled set of multispecies annotations, where we randomly subsample training examples equal to the number of training examples we would have if only considering the species being tested on
2. NetQuilt trained on single-organism annotations, in which we take only rows corresponding to the particular organism being evaluated from the original matrix *S* containing protein similarities among all organisms (for example, training the maxout neural network only on the rows corresponding to human proteins in the block **S** matrix represented in Fig. 1(**B**))
3. Single-species Maxout, in which we take only the IsoRank-score matrix for integrating the single organism’s PPI network with sequence homology information from BLAST, but not including similarities to any other organisms’ proteins (for example, training the maxout neural network only on the **S**_11_ matrix for human proteins represented in Fig. 1(**E**))

These benchmarks allow us to disentangle the effects that the number of training examples and the addition of new features have on performance. In addition to these, we also include deepNF and BLAST (propagating labels from training to test proteins based on sequence similarity as in CAFA [6]). deepNF includes information from STRING network types not used by our models: i.e., the coexpression, cooccurrence, neighborhood, fusion, and database networks. BLAST, like our main multispecies model, uses proteins from all organisms in the set of chosen species to make predictions on the cross-validation test proteins.

### 4.3. Leave-one-species-out validation

The next set of experiments we performed simulate a scenario in which we use the networks of multiple species in order to predict the functions of proteins of an organism with no PPI network available (a reasonably common occurrence for non-model species). An outline of the procedure is shown in Figure 4. We first take a single organism with its annotations left out from training and used as the test set, and leave out the network for that organism. In order to construct the features of the organism for use in the maxout neural network, we first need to obtain interspecies connections between the test organism and all other organisms in the dataset. To do this, we first calculate the sequence similarity between the test organisms’ proteins and all other organisms’ proteins, and run IsoRank in the previously described way, except that we use the identity matrix in place of the PPI network of the left-out organism. We obtain an *n_i_* × *n*_test_ interspecies protein similarity matrix *S*_i,test_ relating each species’ *n*_i_ proteins with the test species’ *n*_test_ proteins. We then perform a one-mode projection, given by 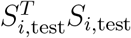, which predicts connections between the nodes of the test species from their shared neighbors (through the IsoRank connections) in other species. Since we have a prediction matrix for every other species in the set besides the test species, we take the element-wise mean of these different matrices to get the predicted network 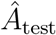. Finally, using this matrix as a proxy for a real PPI network, we run IsoRank on the matrix with itself, combined with its own species’ BLAST connections, to obtain the matrix *S*_test,test_.

## 5. Results

In the following sections, we present the performance of our method in two evaluation settings. The first setting is cross-validation over the annotations of a single species, in which we can compare our method to single-species network-based methods. The second setting is leave-one-species-out (LOSO) evaluation, in which we leave out both a species’ PPI network and its annotations while using the rest of the organisms to train, as outlined in the previous section.

### 5.1. Cross validation over annotations of one species

We present the performance of our method in cross-validation on **human**, **fly**, **mouse** and **E. coli**. We summarize our results using AUPR under micro and macro averaging, accuracy score (Acc) and F1-score (in the same manner as in [12]). The AUPR and F1 scores are computed in a function-centric manner and averaged over all GO terms. Accuracy is computed as the proportion of correctly predicted proteins of the set, where a prediction for a GO term is made if the model outputs a score greater than 0.5. A protein is considered “correctly predicted” if our predictions match the label vector of considered GO terms. We show results separately for the three different branches of Gene Ontology, molecular function (MF), biological process (BP), and cellular component (CC). In Figures 5, 6, and 8, we see that the NetQuilt network trained on model bacteria proteins outperforms the other methods significantly across the three branches of Gene Ontology for E. coli, human and mouse. This can primarily be attributed to the large number examples included in the training set. In addition, the diversity of training examples across multiple species also serves to increase performance, as indicated by the higher performance of the maxout network trained on subsampled sets of annotations from multiple species equal in size to the training set for a single species. For fly, shown in Figure 7, deepNF outperforms our method in the biological process and cellular component branches. We note that deepNF has additional information – the coexpression, cooccurrence, neighborhood, fusion, and database networks – in addition to the experimental network from STRING, while our method incorporates only the experimental network and BLAST connections. The performance of the CAFA BLAST baseline method also performs poorly for fly, which reflects the smaller number and magnitude of BLAST connections between fly and the other organisms (see Supplemental Figure 7 for network and homology comparisons between eukaryotes). This indicates that the homology of the organisms in the set does not give as much information as the other sources of information that deepNF takes into account for the fly protein function prediction task. Since our method also relies on homology information, we expect a corresponding decrease in performance when such information is not as salient to the classification task. We see this effect also in the max-out network trained in the subsampled setting, where homology information from the proteins of other organisms is included in the training data at the expense of other proteins in the fly network.

**Figure 6:**
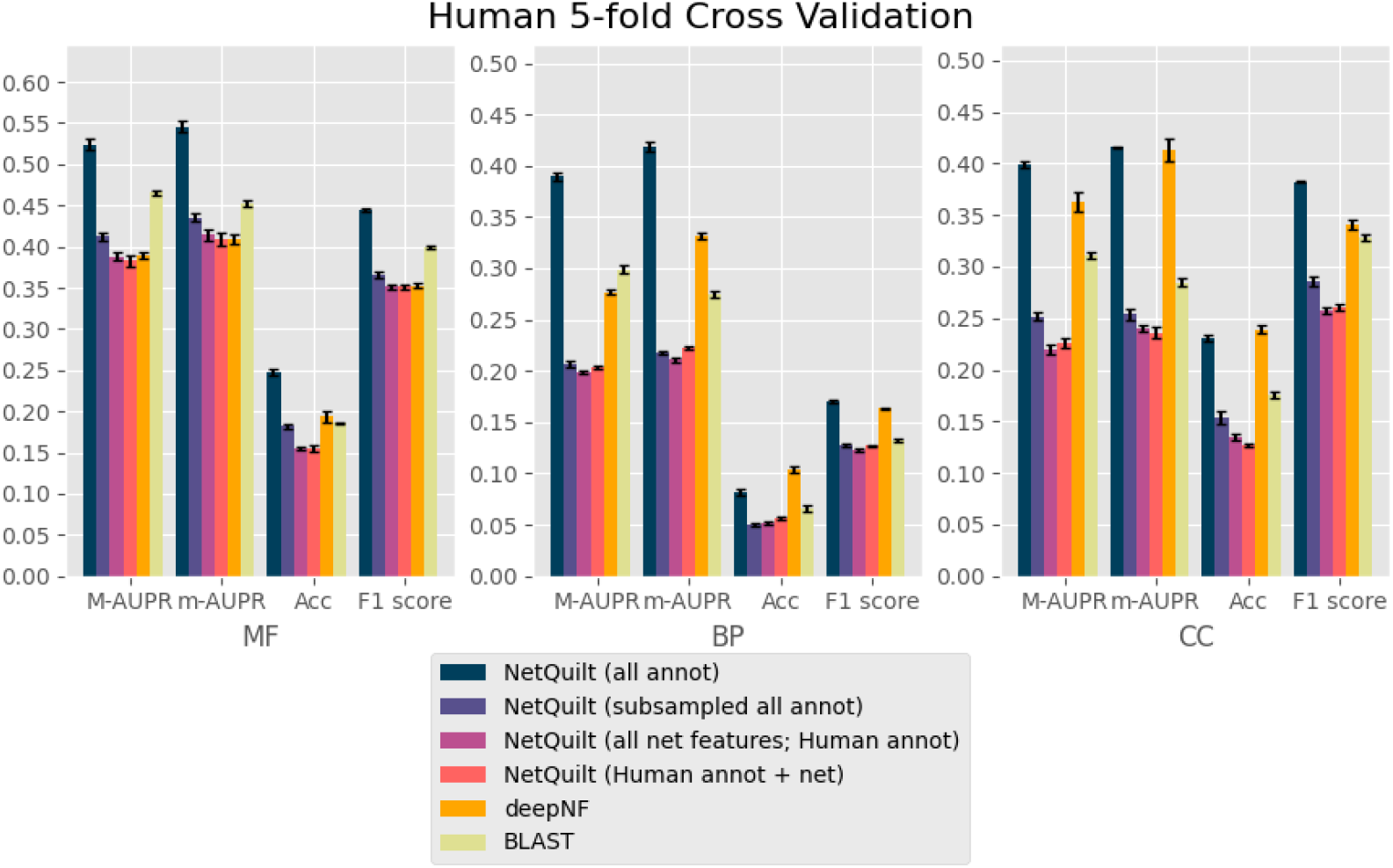
Performance comparison of NetQuilt method with baselines. Methods shown: NetQuilt trained on model eukaryote annotations; NetQuilt trained on subsampled model eukaryote annotations; NetQuilt trained only on human examples; single-species NetQuilt (taking only the human IsoRank matrix and annotations); deepNF (single-species, but integrating 6 STRING network types); and CAFA BLAST annotation transfer method using all selected eukaryote annotations.

**Figure 7:**
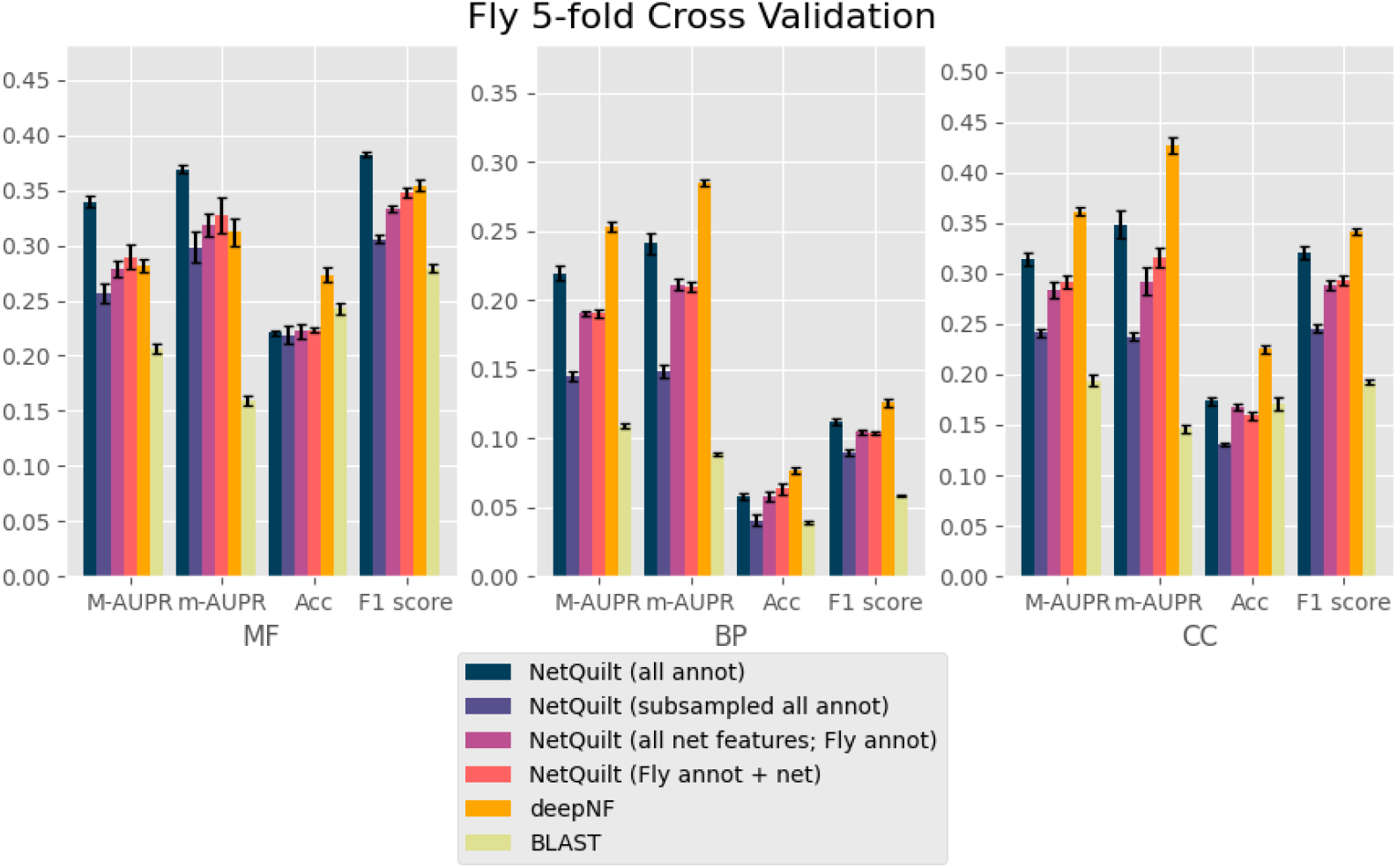
Performance comparison of NetQuilt method with baselines. Methods shown: NetQuilt trained on model eukaryote annotations; NetQuilt trained on subsampled model eukaryote annotations; NetQuilt trained only on D. melanogaster examples; single-species NetQuilt (taking only the fly IsoRank matrix and annotations); deepNF (single-species, but integrating 6 STRING network types); and CAFA BLAST annotation transfer method using all selected eukaryote annotations.

**Figure 8:**
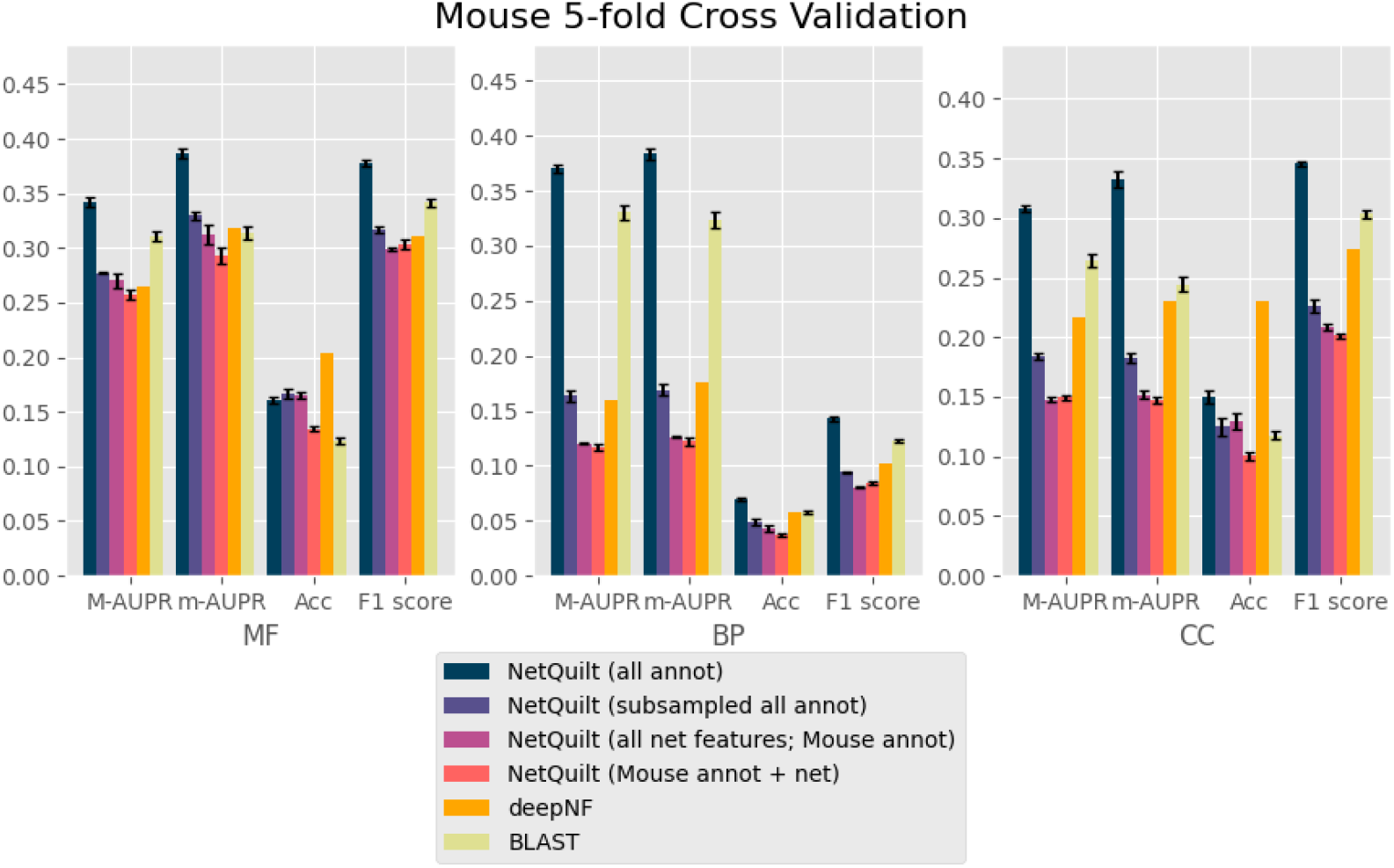
Performance comparison of NetQuilt method with baselines. Methods shown: NetQuilt trained on model eukaryote annotations; NetQuilt trained on subsampled model eukaryote annotations; NetQuilt trained only on Mus musculus examples; single-species NetQuilt (taking only the mouse IsoRank matrix and annotations); deepNF (single-species, but integrating 6 STRING network types); and CAFA BLAST annotation transfer method using all selected eukaryote annotations.

For all organisms, NetQuilt trained only on a single species’ annotations performs similarly whether it uses multispecies features or single-species features. For E. coli and human, training on multispecies features gives slightly better performance with regard to the molecular function ontology than training on single-species features. However, for cross-validation on human in the biological process ontology, the multispecies features actually decrease performance. This may be because adding a significantly larger number of features without increasing the number of training examples has limited benefits, with a higher number of parameters needing more samples to train on. On the other hand, both of these baseline models’ performances are comparable to that of deepNF for the molecular function ontology for all of the considered organisms. This suggests that the features based on PPI networks integrated with homology through our method can enable the neural network to have competitive performance even without large numbers of training examples.

### 5.2. Leave-one-species-out validation

In order to explore the performance of our method in a situation in which no PPI interaction network is known for an organism but homology information is present, we present results for E. coli and fly LOSO validation in Figures 9 and 10, and for human and mouse in Supplemental Figures 1 and 2. This setting often describes the case for many newly sequenced species; mass spectrometry or yeast two-hybrid data may not be available for such organisms.

**Figure 9:**
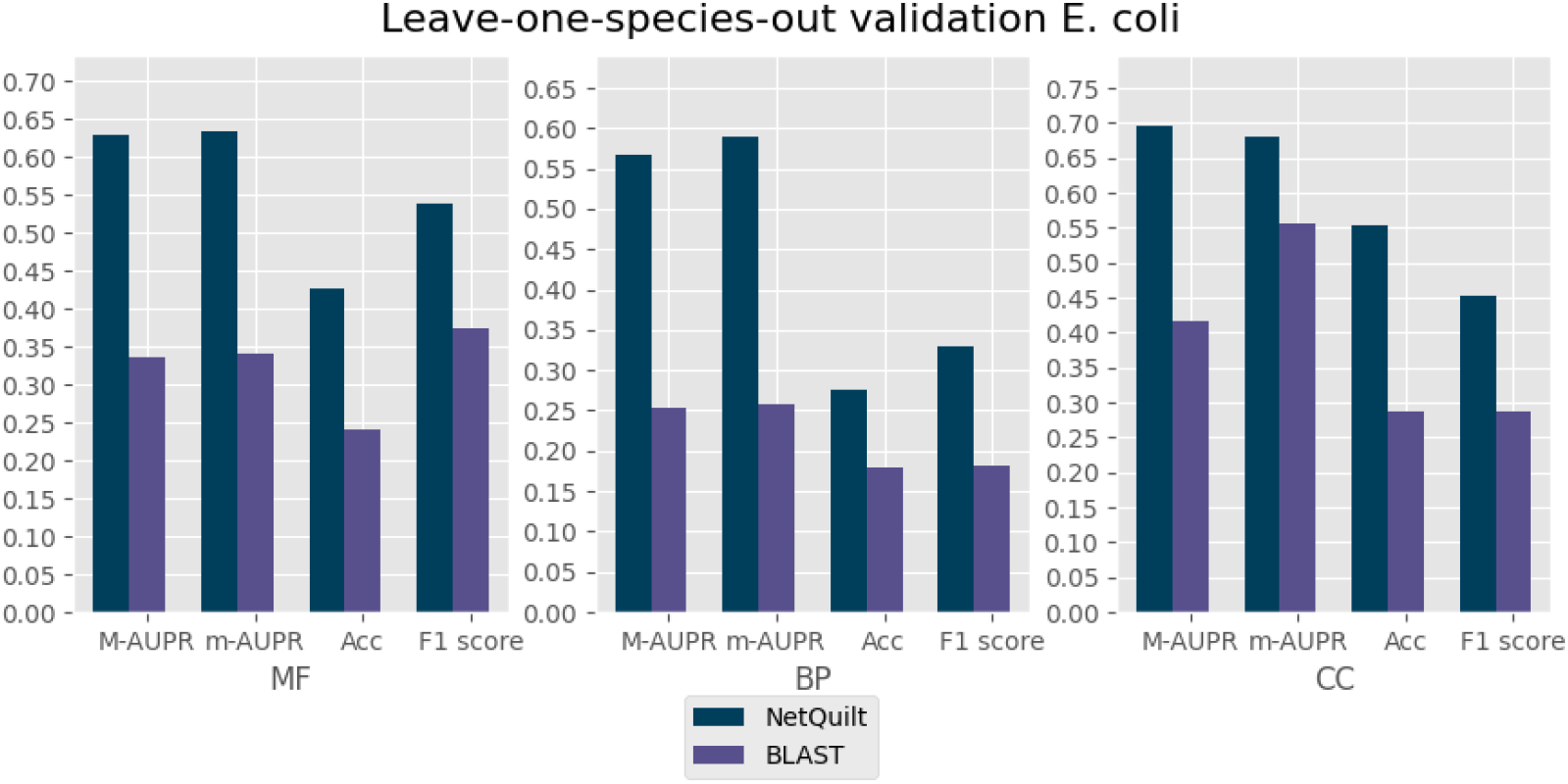
**E. Coli** annotations.

**Figure 10:**
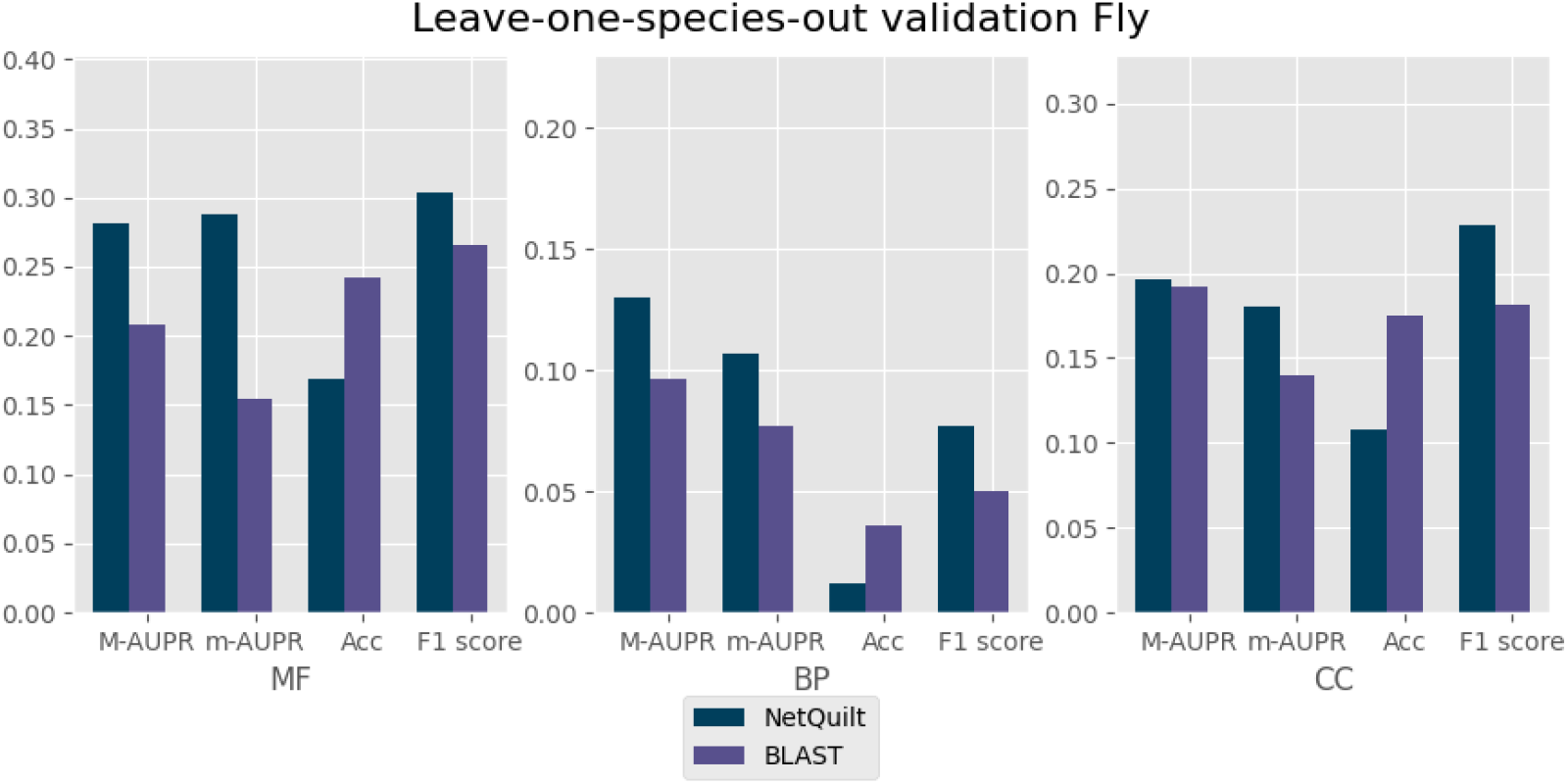
**D. melanogaster** annotations.

For E. coli, we see that our model significantly outperforms the CAFA BLAST labeling method. There are annotations available from all other bacteria, including another well-annotated substrain of E. coli (K-12 substr. W3110; see Figure 3). BLAST can use these presumably useful homologs in transferring annotations to the E. coli K-12 substr. MG1655, our test organism. However, even with this information, our method outperforms BLAST by more than double in the macro-AUPR performance for biological process, and by similarly large margins in the molecular function and cellular component ontologies. For fly, we also see NetQuilt generally outperforming the CAFA BLAST labeling method, though for cellular component, the improvement is not as significant. This shows that our method of integrating multiple species’ PPI networks and their homology link information can be used effectively to annotate proteins for organisms for which neither PPI network nor annotations are available. In particular, it shows that we can outperform strictly homology-based predictions when there is PPI network information available for species related to the organism we want to annotate.

On human and mouse, our model performs approximately as well as the CAFA BLAST-labeling method. The BLAST-labeling method performs much better for these organisms than it does for fly and E. coli. When homology information is highly informative, as is the case in human and mouse, BLAST is difficult to improve upon. However, in cases where homology is not as informative for the annotation task, the complementary PPI data used by our model allows for significant improvements in performance.

## 6. Conclusions and discussion

With the arrival of high-throughput experimental techniques came large PPI network datasets of thousands of organisms. Many function prediction algorithms use PPI information for function prediction using a single species at a time. In order to fully exploit this rich source of information, new protein function prediction algorithms should be designed so that multiple PPI networks can be integrated, along with the most abundant source of protein information: homology. We present here a method that is the first of its kind: a multispecies network-based deep learning method for protein function prediction that effectively integrates PPI network information and homology. The integration of multiple PPI networks is based on IsoRank, a PPI network alignment technique that uses homology to transfer topological similarity scores between nodes of different networks. We use the integrated similarity scores as input to a maxout neural network in order to accurately predict protein function. We demonstrate the superiority of our method in Gene Ontology term prediction to single-species network-based approaches as well as to the homology transfer method from the Critical Assessment of Function Annotation (CAFA) using a cross-validation evaluation.

The multispecies approach enables us not only to produce better predictions in situations involving completing the annotations of a single species using its PPI network, but also to make accurate network-informed predictions on species for which the organism has either an incomplete or an entirely nonexistent PPI network. We show this capability through a leave-one-species-out validation whereby we leave out a species’ network and annotations and train our model on multiple other species, and then evaluate our function predictions on the left-out species. We show that our method can be at least as good as the CAFA homology transfer method in settings in which homology is very informative, and is a great improvement over the CAFA homology transfer method in settings in which homology information is not enough to produce accurate predictions.

This method shows promise for training deep learning models on large multispecies PPI network datasets. In light of the informative representations learned by deep-learning algorithms trained on sequence datasets with millions of training examples, we have a vision of applying deep learning techniques similarly to the millions of nodes in all PPI networks. In future work, we hope to explore principled ways of integrating much larger numbers of PPI networks with homology information for function prediction.

## 7. Appendix

Note that the IsoRank formula in Equation 1 can be interpreted as personalized PageRank for the matrix of the Kronecker product 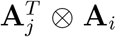 of row-normalized adjacency matrices. This can also be seen as solving the eigenvalue problem for this tensor product graph when *α* =1. Vectorizing the right hand side of equation 1, we have:

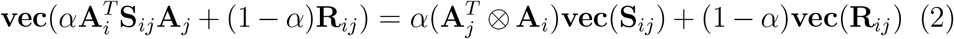

The problem can then be formulated as the following, which we can solve for **S**_*ij*_:

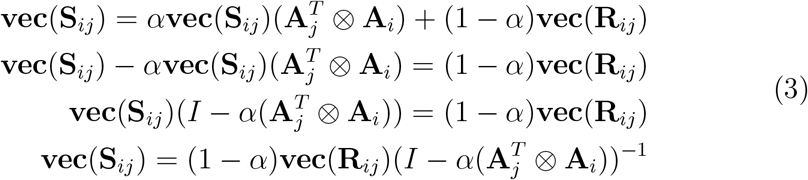

