## Supplementary Material for "NetQuilt: Deep Multispecies Network-based Protein Function Prediction using Homology-informed Network Similarity"

### 1 Network statistics

Table 1: **Network Statistics for CAFA 4 Bacteria Networks.** $\frac{N_c}{N}$ refers to the ratio of largest connected component nodes to the total number of nodes in the graph.

| **Scientific name** | **Taxonomy ID** | **Nodes** | **Edges** | $\mathbf{\frac{N_c}{N}}$ |
|---|---|---|---|---|
| P. aeruginosa | 287 | 6267 | 161805 | 0.703048 |
| E. coli str. K-12 substr. MG1655 | 511145 | 4125 | 66075 | 0.876848 |
| E. coli str. K-12 substr. W3110 | 316407 | 4210 | 61533 | 0.87886 |
| S. enterica subsp. enterica serovar Ty... | 90371 | 4418 | 59412 | 0.774785 |
| M. genitalium G37 | 243273 | 474 | 7361 | 0.814346 |
| B. subtilis subsp. subtilis str. 168 | 224308 | 4181 | 69451 | 0.672088 |
| C. vibrioides CB15 | 190650 | 3721 | 82083 | 0.713518 |
| A. fischeri ES114 | 312309 | 3797 | 74238 | 0.749802 |
| Synechocystis sp. PCC 6803 | 1148 | 3167 | 59624 | 0.843701 |
| P. fluorescens SBW25 | 216595 | 5881 | 245532 | 0.766026 |
| P. fluorescens Pf0-1 | 205922 | 5681 | 225074 | 0.770111 |
| A. vinelandii DJ | 322710 | 4955 | 115245 | 0.699697 |
| S. coelicolor A3(2) | 100226 | 7741 | 224300 | 0.694871 |

Table 2: **Network statistics for Chosen CAFA 4 Eukaryote networks**. $\frac{N_c}{N}$ refers to the ratio of largest connected component nodes to the total number of nodes in the graph.

| **Scientific name** | **Taxonomy ID** | **Nodes** | **Edges** | $\mathbf{\frac{N_c}{N}}$ |
|---|---|---|---|---|
| C. elegans | 6239 | 18181 | 1044754 | 0.66289 |
| D. melanogaster | 7227 | 13046 | 749503 | 0.861873 |
| D. rerio | 7955 | 24681 | 4198604 | 0.93104 |
| H. sapiens | 9606 | 19354 | 2287192 | 0.931745 |
| S. scrofa | 9823 | 21284 | 2892886 | 0.910449 |
| M. musculus | 10090 | 21291 | 2739040 | 0.880607 |
| R. norvegicus | 10116 | 22234 | 3374903 | 0.878205 |

### 2 Leave-one-species-out validation (human and mouse)

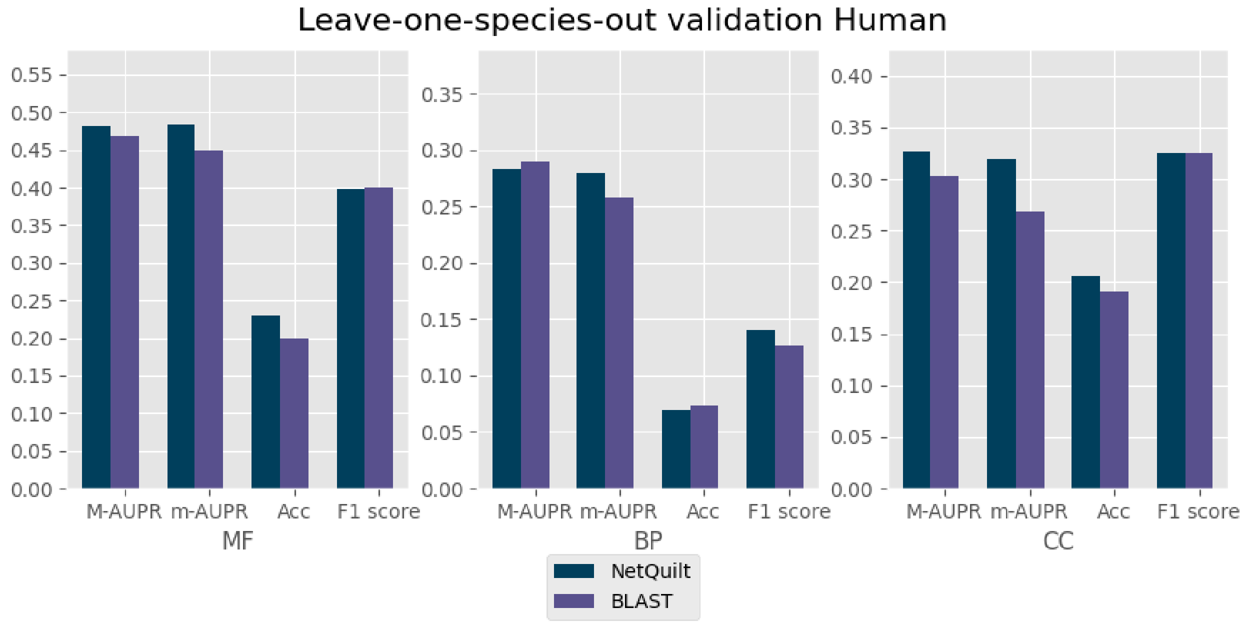


Figure 1: **H. sapiens** annotations.

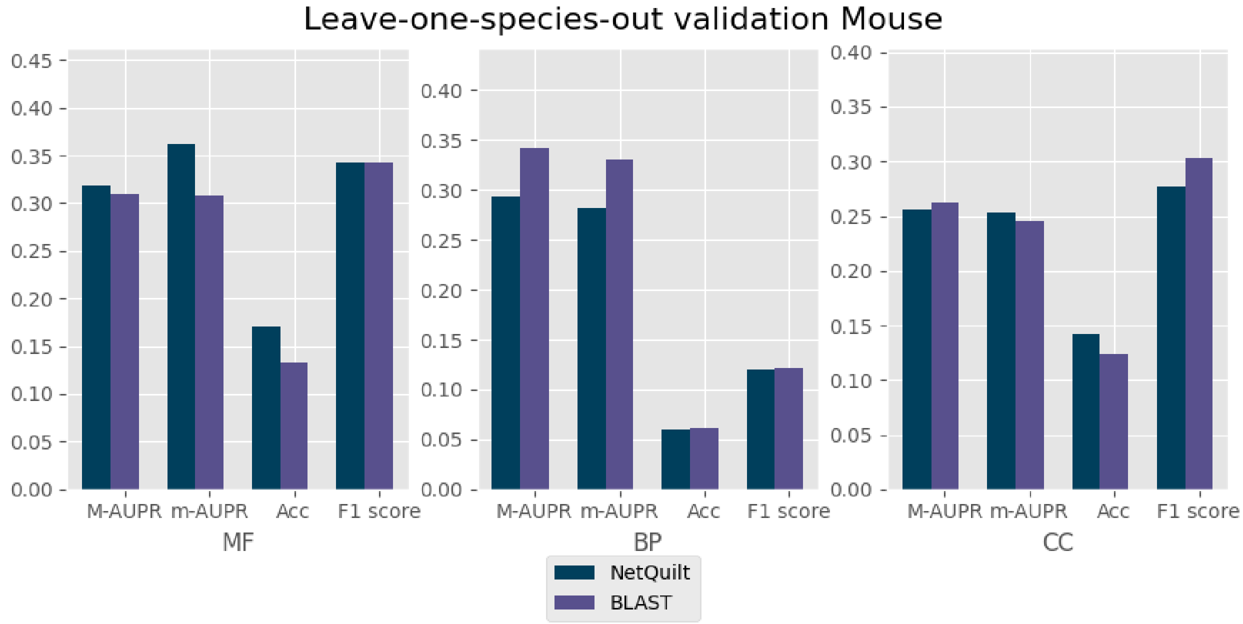


Figure 2: **M. musculus** annotations.

### 3 $\alpha$ search

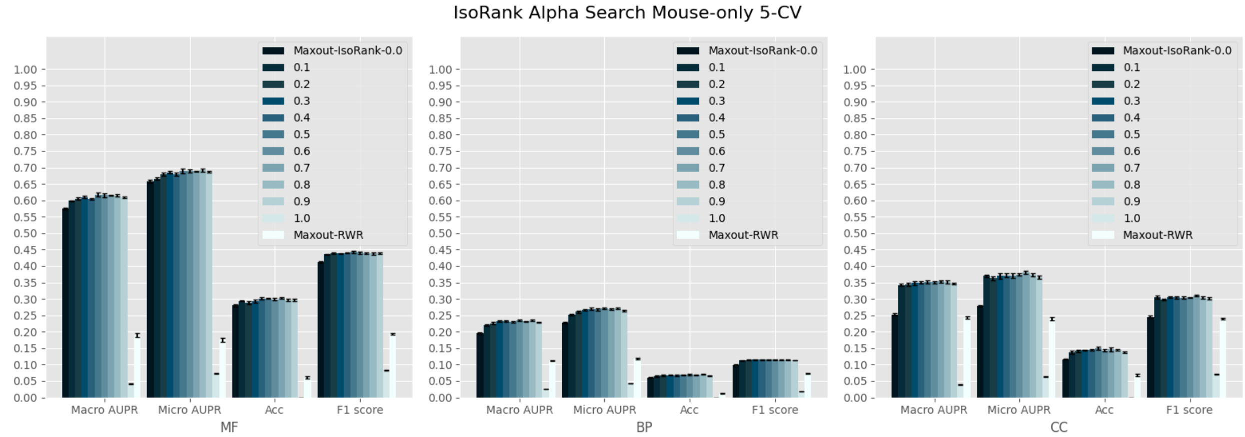


Figure 3: **M. musculus** annotations, cross validation with 5 trials. Maxout-IsoRank-0.0 (and all other bars labeled with numbers) refer to using the mouse IsoRank matrix only, with different values of $\alpha$, as features. Maxout-RWR refers to a maxout neural network trained on a random-walk-with-restarts matrix computed for the mouse PPI network (this method, as well as the IsoRank $\alpha = 1$ setting, creates features with no contribution from homology).

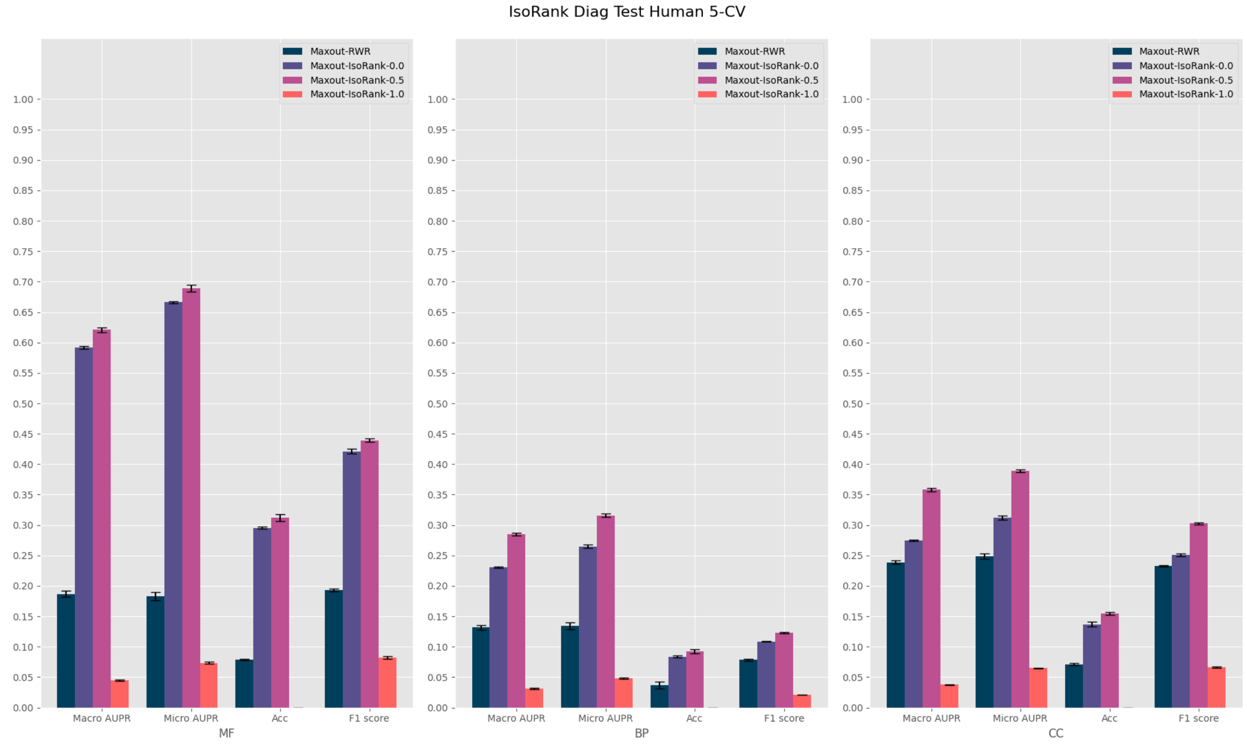


Figure 4: **H. sapiens** annotations and mouse IsoRank matrix only, cross validation with 5 trials. Maxout-RWR refers to a maxout neural network trained on a random-walk-with-restarts matrix computed for the mouse PPI network (this method, as well as the IsoRank $\alpha = 1$ setting, creates features with no contribution from homology). $\alpha = 0.5$ was chosen for all eukaryote experiments as a result of the $\alpha$ search in Supplemental Figure 3.

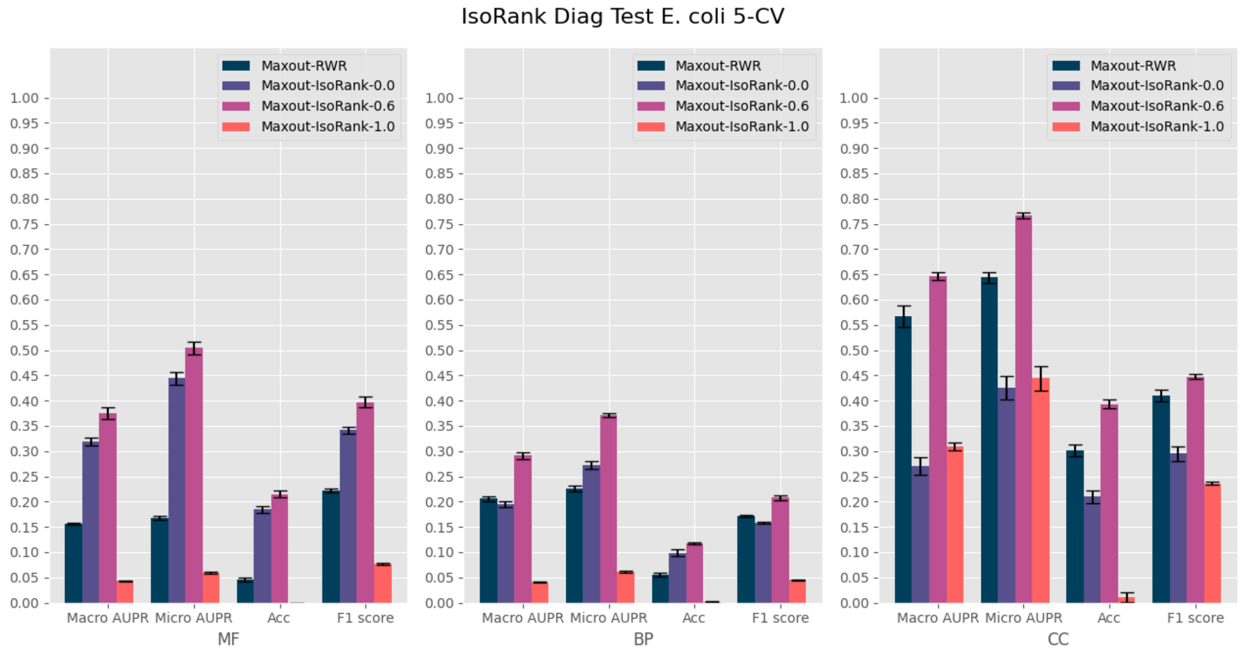


Figure 5: **E. coli** annotations and E. coli IsoRank matrix only, cross validation with 5 trials. Maxout-RWR refers to a maxout neural network trained on a random-walk-with-restarts matrix computed for the mouse PPI network (this method, as well as the IsoRank $\alpha = 1$ setting, creates features with no contribution from homology).

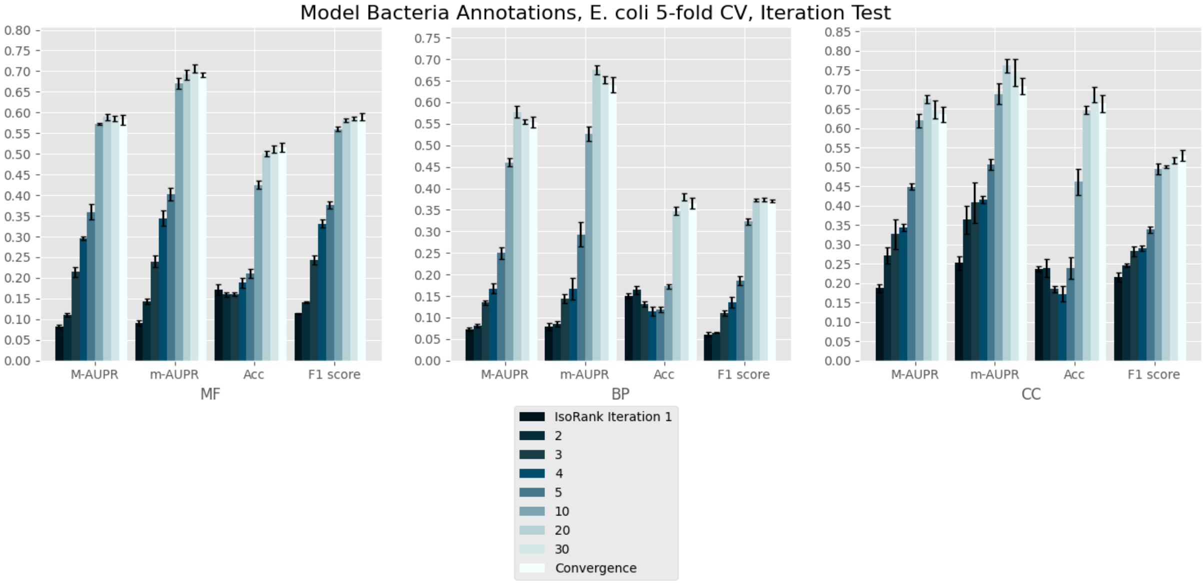


Figure 6: 5 trial CV on E. coli, trained with CAFA bacteria annotations. Test of how performance changes with different iterations of IsoRank.

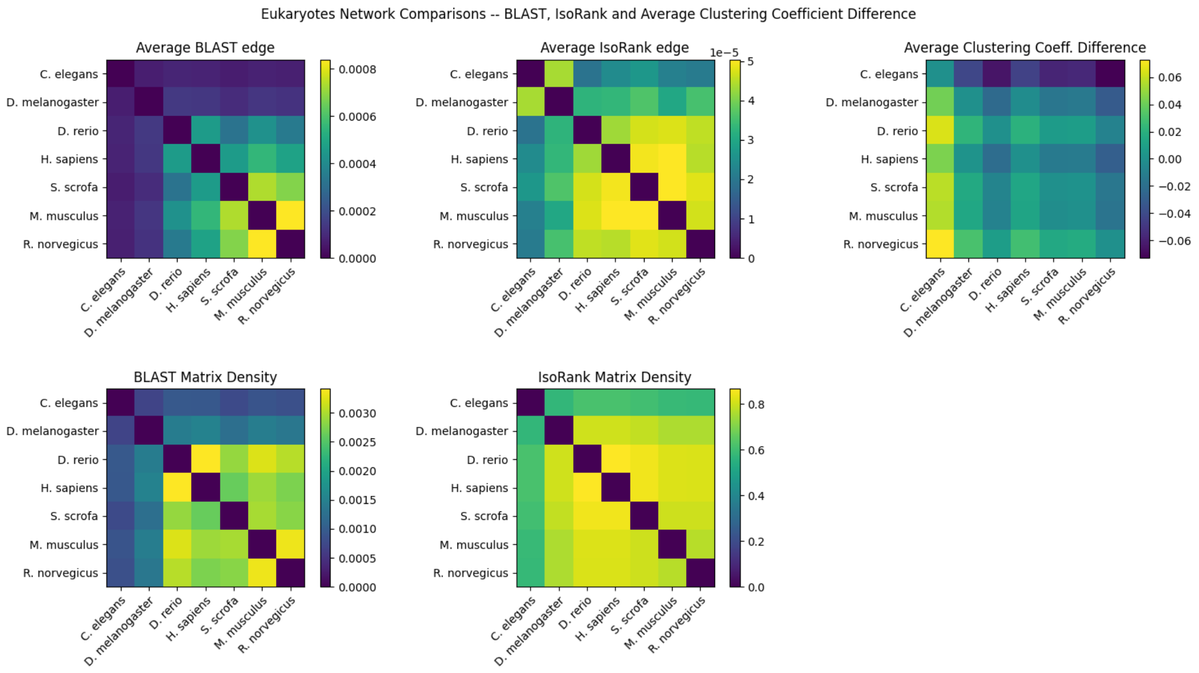


Figure 7: Eukaryote network comparisons. Average BLAST edge refers to the average negative log e-value between the proteins of the two compared organisms, IsoRank edge is the average element of the IsoRank matrix between the two organisms, and the Average Clustering Coeff. Difference is the difference between the average clustering coefficients of the two compared organisms. Since the clustering coefficient difference can be negative, the plot is not symmetric.

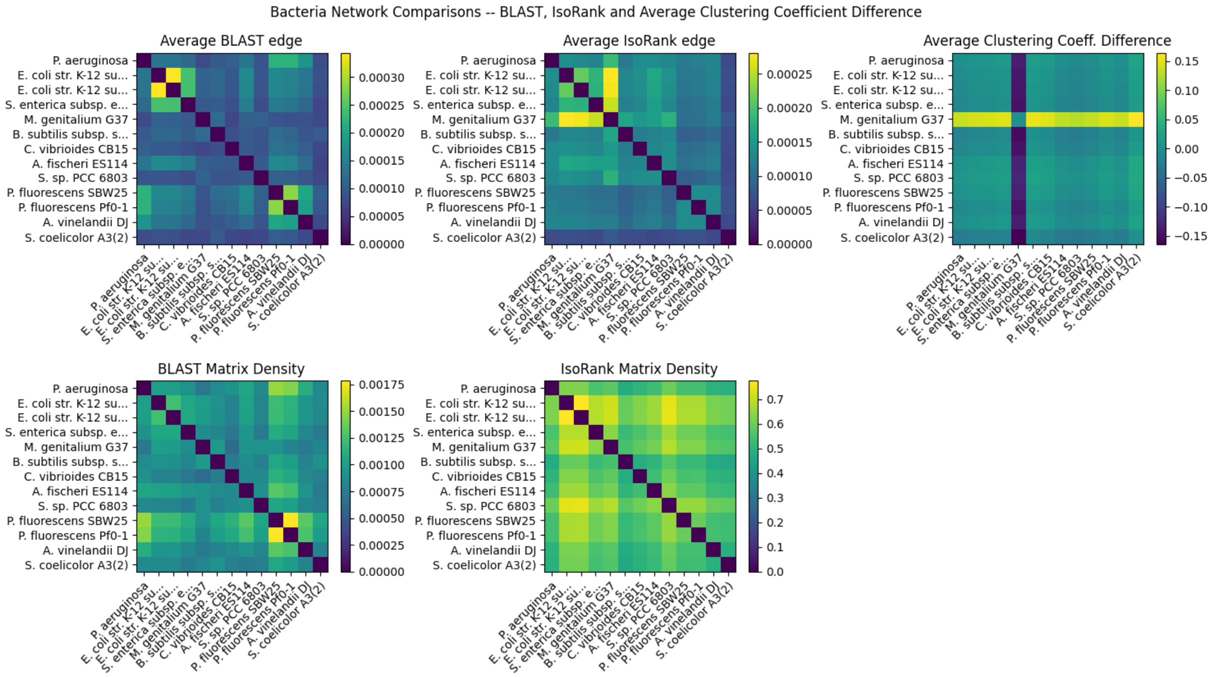


Figure 8: Bacterial network comparisons. Average BLAST edge refers to the average negative log e-value between the proteins of the two compared organisms, IsoRank edge is the average element of the IsoRank matrix between the two organisms, and the Average Clustering Coeff. Difference is the difference between the average clustering coefficients of the two compared organisms. Since the clustering coefficient difference can be negative, the plot is not symmetric.
